# Rapid encoding of temporal sequences discovered in brain dynamics

**DOI:** 10.1101/2020.12.11.421669

**Authors:** L. Bonetti, E. Brattico, F. Carlomagno, G. Donati, J. Cabral, N.T. Haumann, G. Deco, P. Vuust, M.L. Kringelbach

## Abstract

Information encoding has received a wide neuroscientific attention, but the underlying rapid spatiotemporal brain dynamics remain largely unknown. Here, we investigated the rapid brain mechanisms for encoding and prediction of sounds forming a complex temporal sequence. Specifically, we used magnetoencephalography (MEG) to record the brain activity of 68 participants while they listened to a highly structured musical prelude. Advanced analysis of the phase synchronisation and graph theoretical measures showed the rapid transition of brain activity from primary auditory cortex to higher order association areas including insula and superior temporal pole within a whole-brain network, occurring during the first 220 ms of the encoding process. We discovered individual differences, revealing the rapid unfolding of brain network dynamics responsible for the processing of the current sounds and the prediction of the forthcoming events of the sequence. This provides a first glimpse of the general mechanisms underlying pattern encoding in the human brain.

## Introduction

Memory is doubtless one of the most crucial cognitive abilities of humans and animals, necessary to allow species to survive and act^1^. Among its building elements, information encoding plays a fundamental role, allowing individuals to accomplish the crucial goal of learning from experience. In the last decades of neuroscience, much has been done on information encoding, working both with humans and animal models^2–4^. A large part of this research focused on encoding of visual and spatial stimuli^5,6^, whereas a conspicuous amount of studies explored auditory processes related to early and even pre-attentive elaboration of standard and deviant sounds inserted in elementary sequences^7–9^.

The former approach led to several examples of visual encoding and recognition studies which employed faces as main stimuli. Along this line of research, neuroscientists have revealed the primary role of fusiform gyrus in face encoding and recognition and shown the progressive dynamics of different face features elaboration^10–12^.

Conversely, regarding auditory research a large number of studies deeply explored the early and pre-attentive processing of sounds and simple temporal sequences. For instance, this research highlighted several automatic event related potentials/fields (ERP/F) to standard and deviant sounds such as the well-known N100 and mismatch negativity (MMN). Indeed, it has also shown that N100 and MMN were modulated by the stimuli characteristics, suggesting a primitive intelligence and memory of the primary auditory cortex^7–9^. However, this research mainly involved passive listening paradigms and studied the automatic brain responses to sound stimulations, while it did not directly explain how the sounds were actually encoded. A more general approach on sound processing investigation reported the activation of auditory cortex brain regions such as Heschl’s and superior temporal gyri in response to acoustic stimuli varying in temporal and spectral features^13,14^ as well as a contribution coming from inferior frontal gyrus, cingulo-insular cortex, mediotemporal limbic lobe, basal ganglia, supplementary motor area and posterior orbitofrontal cortex^15,16^.

The encoding of temporal sequences has also been investigated by applying more complex paradigms, especially within the domains of language and algebraic representation. For instance, the analysis of the brain activity underlying the attentive processing of sound sequences comprising regular subgroups of items returned not only the well-known automatic mismatch response (MMN) to deviant sounds, but also a late surprising-elicited P3b component when the coherence of the sequence was broken^17,18^. These studies have suggested that the brain actively chunked the different pieces of information that were presented. Others studies demonstrated that the ordinal arrangement of a series of items is encoded by the brain and represents a very important feature of the learning process of the presented stimuli. Such brain mechanisms have been detected both in humans and monkeys showing a neural code for ordinal numbers and items ordinal representation within the intraparietal and dorsolateral prefrontal cortices^19,20^. Further research employing animal models and temporal sequences characterized by constant algebraic patterns suggested that naive non-human primates were able to represent the abstract numerical and algebraic patterns of sequences, understanding a higher level of abstract regularities than elementary sequences with sensorial deviations^21,22^.

To review the decades of sequence encoding findings within cognitive sciences and neuroscience, a recent key paper by Dehaene, Meyniel, Wacongne, Wang and Pallier^23^ proposed a taxonomy of the encoding mechanisms for temporal sequences, elaborating five distinct categories. They refer to transition and timing knowledge between subsequent items *(i)*, chunking of contiguous items of the sequence *(ii)*, ordinal knowledge of which item comes first *(iii)*, algebraic patterns capturing complex regularities within a sequence *(iv)*, nested tree structures based on abstract symbolic rules *(v)*. However, even though Dehaene et al.^23^ provided a detailed description of the brain responses to sequences evolving over time, they did not reveal the temporal dynamics of the subsequent brain processes required for the encoding and prediction of sounds forming a complex temporal sequence. Moreover, as explicitly stated in their work, Dehaene et al.^23^ highlighted the urgency to unravel which is the specific contribution of cortical and subcortical brain networks to the extraction of the sequence structure and the prediction of its forthcoming elements.

Thus, here we used music, which is the human art that mainly acquires meaning through the logical combination of sounds extended over time^24^, to investigate the rapid whole-brain networks underlying the encoding and prediction of items forming a complex temporal sequence. Understanding such topic is a key step to discover the general neural mechanisms underlying pattern encoding and thus unveil how the information becomes meaningful for the human brain.

## Results

### Experimental design and data analysis overview

In our study, we aimed to investigate the fine-grained spatiotemporal dynamics of the brain during the encoding and prediction of sounds forming a complex temporal sequence. To this aim, we used MEG to record the brain functioning of 68 participants while listening to a musical instrumental digital interface (MIDI) version of the full prelude in C minor BWV 847 composed by Johann Sebastian Bach.

As described in the Methods and depicted in **Figure 1A**, participants were requested to attentively listen to the music, trying to memorize its structure and sounds as much as they could. The analysis pipeline employed in our study is depicted in **Figure 1** (and described in details in the Methods). Our results on brain functioning underlying sound encoding have been then organised as follows: 1) sensor space and beamformed source localised activity, 2) static source localised connectivity, 3) dynamic source localised connectivity and 4) dynamic source localised connectivity in subsamples characterized by different levels of general (GWM) and auditory working memory (AWM) and musical expertise.

**Figure 1.**
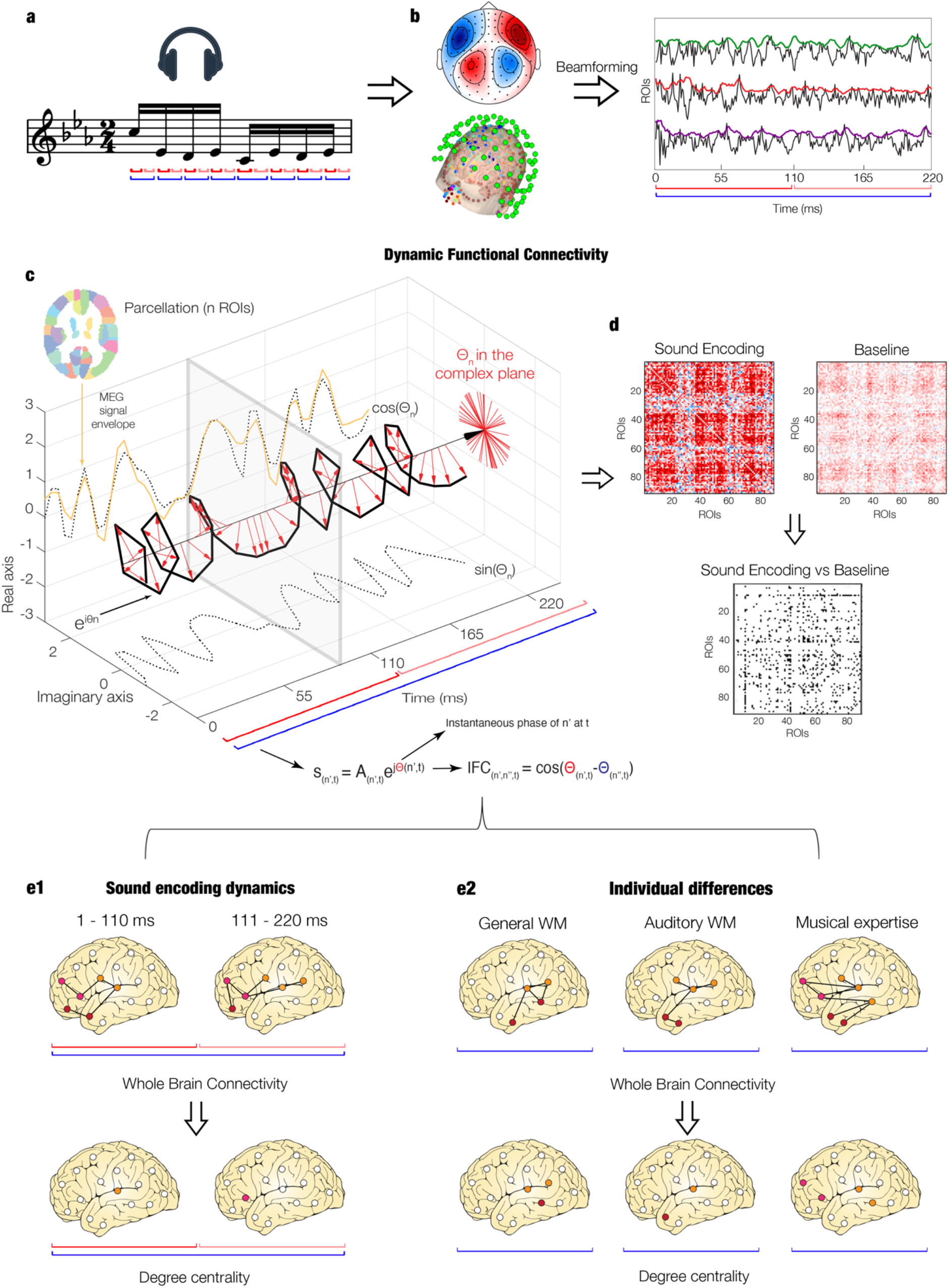
Overview of the analysis pipeline. **a** – Participants were requested to attentively listen to a whole musical piece trying to remember it as much as possible. Subsequent analyses focused on two main time-windows of the sound encoding process, as illustrated by the red and blue lines **b** - MEG data during musical encoding has been collected, pre-processed and beamformed into source space within the 90 non-cerebellar brain regions of the AAL parcellation. Then, we calculated the envelope of the time-course of each brain region (top right). **c** – After computing analysis on brain activity and SFC, we investigated the DFC. To this aim, we computed the Hilbert transform of the envelope of each brain region and estimated the phase synchronization by calculating the cosine similarity between the instantaneous phases of each couple of brain regions. **d** – We obtained IFC matrices for both sound encoding task and resting state (used as baseline). Afterwards, we contrasted the task matrices vs the average of the baseline matrices to isolate the brain activity specifically associated to the sound encoding brain processes over time. **e1** – We used the whole-brain connectivity measures to derive the significant centrality of the brain regions within the whole-brain network. This was done for the two main time-windows of sound encoding (1 – 110 ms and 111 – 220 ms). **e2** – Similarly, contrasting participants who were grouped according to their level of WM and musical expertise, we computed the significant centrality of brain regions associated to those different skills.

First, we detected the brain activity in MEG sensor space using univariate tests and Monte Carlo simulations (MCS). Then, we reconstructed the sources of the brain signal using a beamforming algorithm (**Figure 1B**). Second, we computed the static functional connectivity (SFC) by calculating Pearson’s correlations between the envelope of each couple of brain areas. Third, we computed dynamic functional connectivity (DFC) using the instantaneous phase obtained from Hilbert transform for each time-point of the brain areas timeseries (**Figure 1C**). After contrasting the brain connectivity patterns for sound encoding vs resting state (**Figure 1D**), we computed the DFC for two short time-windows (1 – 110 ms, 111 – 220 ms), as depicted in **Figure 1E1**. Fourth, we analysed the DFC in subsamples of individuals characterized by different levels of GWM, AWM and musical expertise (**Figure 1E2**). DFC analysis consisted of detecting the whole-brain connectivity patterns and then the significant centrality of specific brain areas within the whole-brain network.

### Event related fields and power spectra analysis

At first, as depicted in **Figure 2A** and **2C** to assess the quality of the data, we analysed the ERF associated to the processing of the tones. To this aim, after carrying on standard pre-processing steps (see Methods for details), we epoched the data in correspondence to each musical tone (trials), we averaged the trials and combined planar gradiometers by mean root square. Then, we calculated a t-test for each time-point in the time-range 0.050 – 0.200 seconds and for each gradiometer channel contrasting the task vs its own baseline. Finally, we performed cluster-based MCS to detect significant clusters (t-test threshold = 1.0e-16, MCS threshold = .001). Results showed two significant clusters in the time-range 0.053 – 0.160 seconds. Specifically, we observed a larger cluster in the right hemisphere (cluster size: 81, *p* < .001) and a smaller one in the left (cluster size: 40, *p* < .001). These results are reported in details in **Table ST1**. Then, we reconstructed the neural sources of the signal by applying a beamforming approach and calculating a general linear model (GLM) for each source voxel and each time-point within the significant time-range emerged by MCS on sensor data (0.053 – 0.160 seconds). Significant clusters were assessed by a cluster-based permutation test. **Figure 2D** shows that the activity was mainly localized within primary auditory cortex and insula. Complete results are reported in **Table ST2**. Finally, to convey our subsequent functional connectivity analysis to a specific frequency band, we calculated the power spectra associated to the task by complex Morlet wavelet transform (from 1 to 60 Hz with 1-Hz intervals) to define which frequencies were involved in the sound processing. We then calculated t-tests for each frequency and time-point within the range 0.050 – 0.200 seconds and the averaged power spectra of the baseline. Binarized values (threshold = 1.0e-18) were submitted to a two-dimensional (two-D) MCS (threshold = .001). As reported in **Figure 2B**, the analysis returned a significant cluster (size: 69, *p* < .001) for frequency range: 3 – 5 Hz within the time-range: 0.053 – 0.200 seconds.

**Figure 2.**
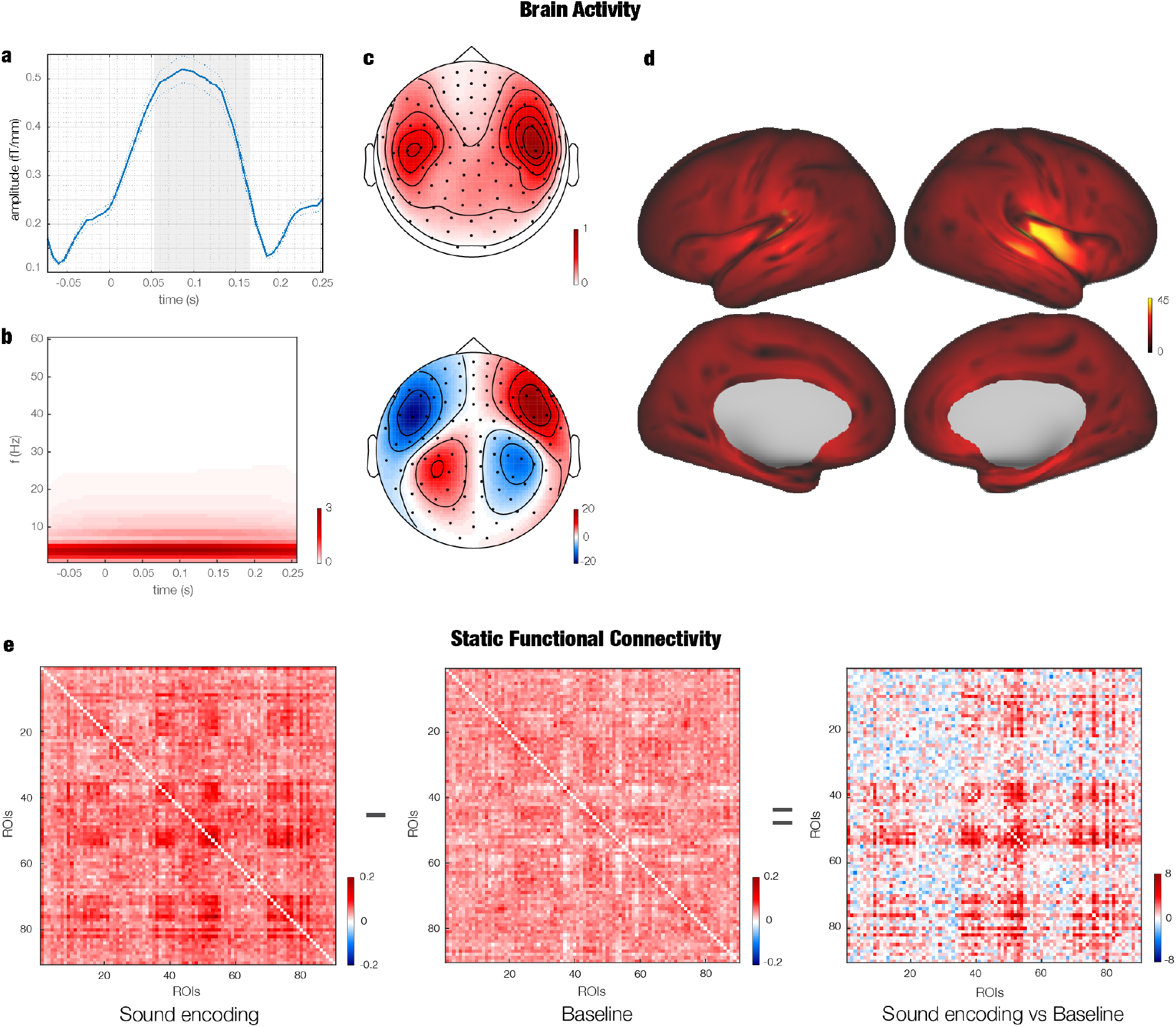
N100 component. **a** – Waveform depicting the N100 component of the ERF. The plot shows the brain activity averaged over the significant gradiometer channels emerged by MCS. The grey area illustrates the significant time-window. Thinner lines depict standard errors. **b** – Power spectra depiction over time for all MEG channels. **c** – Gradiometers (top) and magnetometers (bottom) topoplots of the N100 component amplitude computed over the significant timewindow emerged by MCS. Values correspond to neural signal in fT/mm for gradiometers and fT for magnetometers. **d** – Neural sources of the N100 component (t-values from group level analysis). **e** – SFC estimated by computing Pearson’s correlations between the envelope of each couple of AAL brain regions. The left matrix refers to sound encoding task, while the middle one to resting state (used as baseline). Finally, the right matrix depicts the *t*-values emerged by contrasting task vs baseline.

### Static functional connectivity

We constrained the reconstructed sources of the MEG signal from the 3559 voxels outputted by the beamforming algorithm to the 90 non-cerebellar parcels of automated anatomical labelling (AAL) parcellation and corrected for source leakage. Then, we calculated the SFC by computing Pearson’s correlations between the envelope of the timeseries of each couple of brain areas. This procedure was carried out for both task and resting state. Then, we contrasted the SFC patterns of the sound encoding vs resting state to identify the brain connectivity properly associated to the experimental task. Finally, we used a degree MCS (MCS threshold = .001) for assessing which brain areas were significantly central within the whole brain network during the sound encoding process (see Methods for details).

As depicted in **Figure 2E** and **Figure SF2**, the analysis returned a significant centrality of the following brain regions: left precentral (*p* = 3.5e-05), Rolandic operculum (*p* < 1.0e-07), caudate (*p* < 1.0e-07), putamen (*p* = 4.4e-06), thalamus (*p* < 1.0e-07), Heschl’s gyrus (*p* = 3.0e-05), temporal superior (*p* < 1.0e-07), right temporal pole middle (*p* = 6.6e-06), temporal pole superior (*p* < 1.0e-07), temporal superior (*p* = 1.8e-05), Heschl’s gyurs (*p* < 1.0e-07), thalamus (*p* < 1.0e-07), pallidum (*p* < 1.0e-07), putamen (*p* < 1.0e-07), amygdala (*p* < 1.0e-07), hippocampus (*p* < 1.0e-07), insula (*p* < 1.0e-07), frontal medial orbital cortex (*p* = 6.5-e-04), subgenual (*p* < 1.0e-07), Rolandic operculum (*p* < 1.0e-07), frontal inferior operculum (*p* = 8.2e-05).

### Dynamic functional connectivity

To unravel the dynamics of the functional connectivity during sound encoding, we calculated the instantaneous phase of the signal envelope of matrix by applying Hilbert transform and we estimated the phase synchronisation between each couple of brain areas by computing the cosine of the difference of those instantaneous phases (**Figure 1C**). As done for SFC analysis, to estimate the instantaneous connectivity associated to the task, we contrasted the sound encoding vs rest DFC matrices using Wilcoxon sing-rank test. Then, as described above, a degree MCS assessed the significantly central brain regions within the brain network. Additionally, in this case we were able to describe and contrast using MCS the functional connectivity for two subsequent time-windows (1 – 110 ms and 111 – 220 ms). As illustrated in **Figure 3** and **Figure SF3,** the results highlighted that right Rolandic operculum (*p* < 1.0e-07) and Heschl’s gyrus (*p* = 2.2e-06) were more central within the first 110ms, while right insula (*p* = 2.2e-06) and superior temporal pole (*p* = 2.2e-06) in the second time-window.

**Figure 3.**
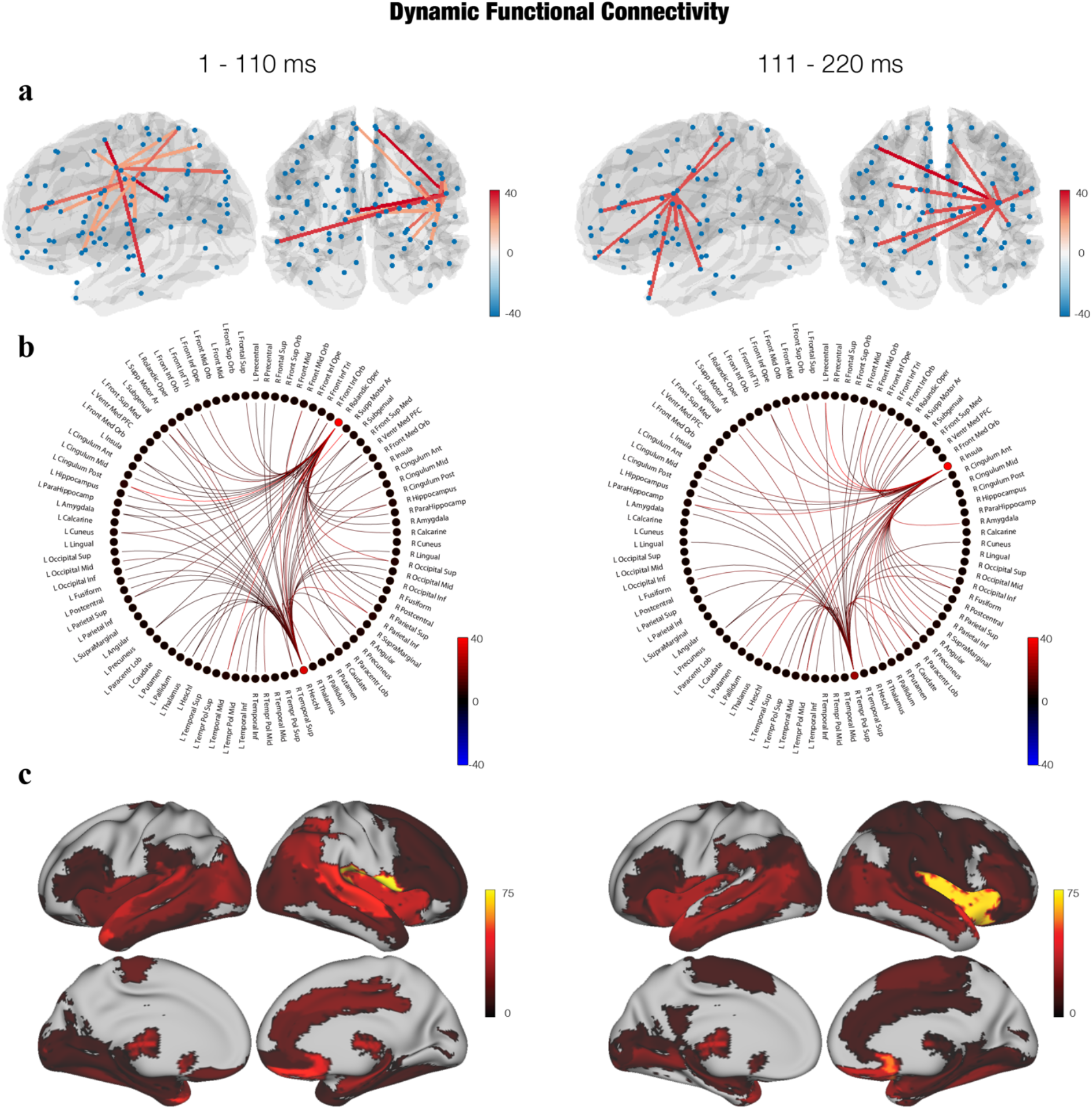
Dynamic functional connectivity. **a** – Connectivity depicted within a brain template showing the strongest connections between the significantly central brain regions and the rest of the brain during two main time-windows (1 – 110 ms on the left and 111 – 220 ms on the right). For each couple of brain templates, the left one is a refiguration from the left hemisphere, while the right one a posterior depiction. **b** – Schemaball showing the strongest connections between the significantly central brain regions and the rest of the brain during the two time-windows. **c** – Significantly central brain regions within the whole brain network during sound encoding depicted in the two time-windows. In all depictions, the colorbar values refer to the temporal extent (in ms) of the brain regions significance.

### Dynamic functional connectivity and individual differences

In conclusion, we aimed to assess whether the neural networks activated during sound encoding differed across participants grouped according to WM abilities (general, assessed by Wechsler Adult Intelligence Scale-IV (WAIS-IV), and auditory, assessed by Musical Earing Test (MET)) and musical expertise. To highlight the differences, we selected for each WM test two groups formed by participants that were at least one standard deviation (SD) apart from each other (see Methods for details). With regards to musicianship, we selected only non-musicians and musicians who had a formal training for at least 10 years. Then, for each of the three variables, we calculated independent contrasts between the two groups using MCS.

**Figure 4** shows that higher GWM was associated to higher centrality of right Rolandic operculum (*p* = 1.2e-05) and lower GWM to left (*p* = 8.4e-05) and right putamen (*p* = 8.4e-05). With regards to AWM skills, the best participants reported higher centrality of right (*p* < 1.0e-07) and left insula (*p* = 6.6e-06), left frontal middle orbital cortex (*p* = 1.1e-06), right temporal middle gyrus (*p* = 6.6e-06) while worst ones had a higher centrality of right occipital inferior (*p* < 1.0e-07), occipital superior (*p* = 6.4e-05) and frontal medial orbital cortex (*p* = 6.4e-05). Finally, musicians exhibited higher centrality of right insula (*p* < 1.0e-07), subgenual cortex (*p* = 5.7e-05), left supplementary motor area (*p* = 1.1e-06), while non-musicians of right caudate (*p* < 1.0e-07) and occipital inferior (*p* < 1.0e-07), as illustrated in **Figure 4**. Additional information about WM, analysis methods and sound encoding brain networks related to all participants are provided in the Methods section and in Supplementary Materials (**Figure SF1**).

**Figure 4.**
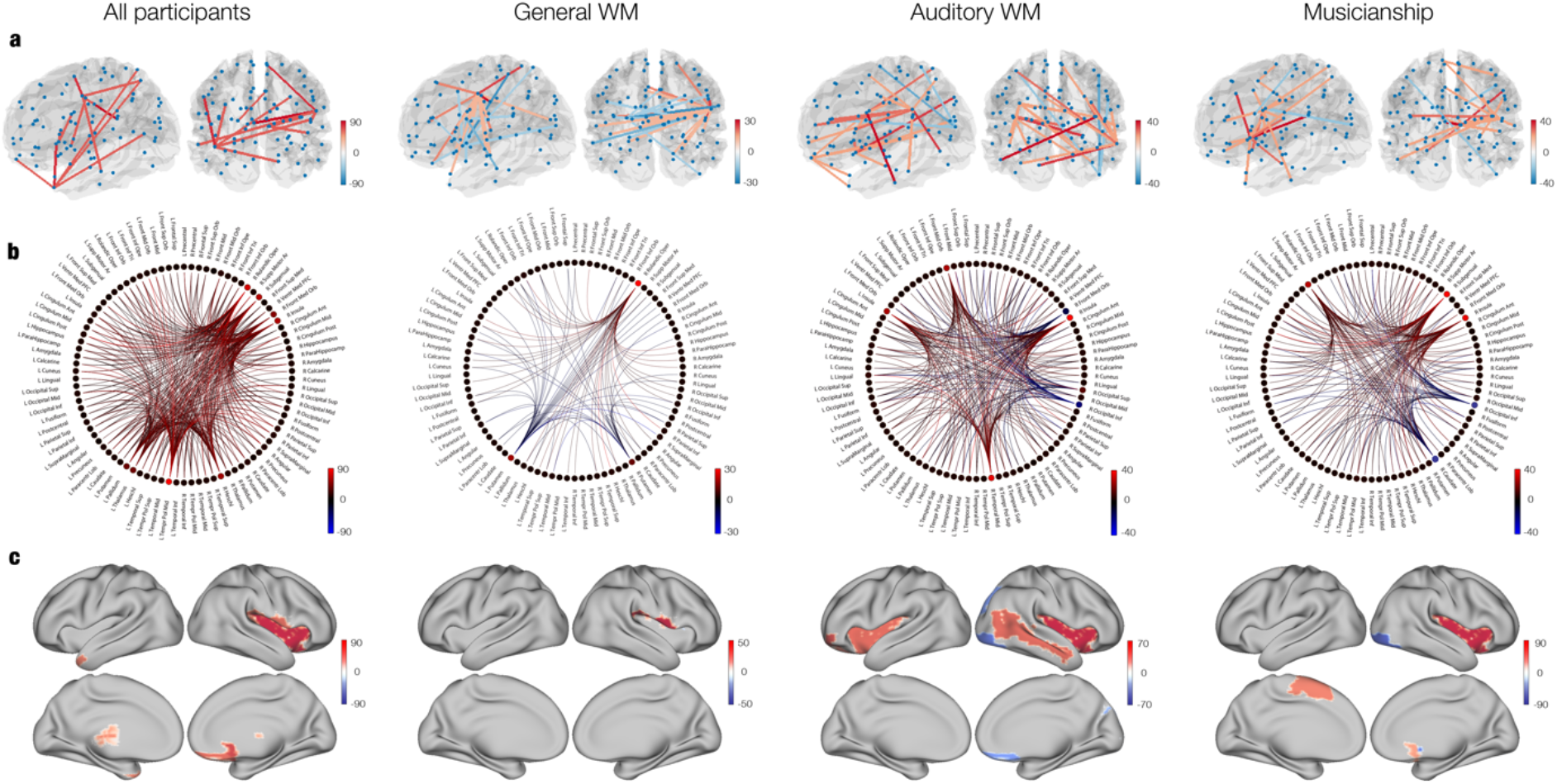
General and auditory WM, musical expertise and brain dynamics. **a** – Depiction within brain templates of the connectivity between the significantly different brain regions emerged contrasting participants’ groups and the rest of the brain. Left column refers to all participants to have an immediate visual comparison, middle-left column represents participants grouped for general WM abilities, middle right relates to auditory WM abilities, while the right column shows the contrast between musicians and non-musicians. For each couple of brain templates, the left one is a posterior refiguration from the left hemisphere, while the right one a posterior representation. **b** – Schemaball depiction of connectivity between significantly different brain regions emerged contrasting participants’ groups and the rest of the brain. **c** – Significant differences of brain regions centrality for contrasts among the different groups of participants. All colorbars depict high scorers and musicians with red shades and low scorers and non-musicians with blue shades. Values show the temporal extent (in ms) of the significant differences.

## Discussion

In this study, we investigated the rapid spatiotemporal brain mechanisms for encoding and prediction of sounds forming a complex temporal sequence. Indeed, we highlighted the specific contribution of cortical and subcortical areas to the brain networks implicated in processing the current item of the sequence and predicting the forthcoming ones. Notably, investigating the brain functioning within the first 220 ms after sounds onset, we detected the brain mechanisms that are presumably responsible to make temporal information available to human awareness^25^.

We detected significant activation and centrality, primarily in the right hemisphere, of several brain regions linked to memory, attentional and auditory processes such as primary auditory cortex, frontal operculum, basal ganglia, insula and hippocampus. Additional analysis employing phase synchronization and therefore dynamical changes over time of the connectivity patterns highlighted stronger centrality of auditory cortex regions such as right Heschl’s and superior temporal gyri as well as frontal operculum within the first 110 ms of the processing of each sound. Conversely, the second time-window that we defined (110 – 220 ms) showed a higher centrality of right insula and superior temporal pole, suggesting the potential role of such structures within the whole-brain network to predict the forthcoming elements of our complex temporal sequence. In conclusion, we presented results connecting individual differences among WM skills and the brain network underlying sound encoding. Specifically, stronger auditory WM skills and musical expertise were linked to higher centrality of subgenual cortex, insula and supplementary motor area, while higher general WM abilities were connected to stronger centrality of right frontal operculum.

### Brain activity and static functional connectivity

Coherently with our results, primary auditory cortex has been widely shown responsible for processing of sound stimulations by a number of well-known studies^26,27^ Additionally, we detected the strongest activity within the right hemisphere, another result largely described and replicated by previous literature^28^. Interestingly, the study of connectivity allowed us to detect several significantly central brain regions that are not directly implicated in auditory processing. Notably, this result suggested the relevance to conduct both activity and connectivity analyses to obtain a complete picture of the sound encoding processes. Studying connectivity, the significant centrality was observed for a number of brain regions such as hippocampus, an area repeatedly connected to memory encoding^29–31^ and frontal operculum, a brain region that has been linked to linguistic production and processing^32,33^. Another key central area that we detected was insula, whose involvement could be related to the salience of the stimuli to be encoded, coherently with studies that showed the role of insula in processing stimulation salience^34,35^. Basal ganglia also played a role in the sound encoding brain network. These subcortical regions have been shown important for several different tasks and are likely most known for their involvement in motor activities and associative learning^36,37^ Since in this study participants were actively attempting to memorize the musical piece, basal ganglia centrality within the brain may be interpreted as a sign of the learning process occurring while listening to the piece.

### Dynamic functional connectivity of sound encoding

A further relevant result that we achieved comes from the study of functional connectivity rapid dynamics that allowed us to identify two main time-windows of sound encoding brain processes. This procedure returned two similar networks of brain regions that were however differentiated by a diverse centrality of few areas. Primary auditory cortex was central mainly in the early time window whereas its connectivity was less marked in the later time interval which was more distant from the sound onset. This result relates to previous literature highlighting its role for the first sensorial processing of upcoming auditory stimuli^38^. More surprisingly, the same result regarded frontal operculum which could be as well important for the brain network associated to the first processes of sound encoding. On the contrary, right insula was predominantly central within the second time-window. In accordance to what we described above, insula may relate to salience of the stimuli and in this case could have played an important role for achieving a more fine-grained categorization of the upcoming sounds. Once again, this interpretation would be coherent with previous studies highlighting insula’s role in salience appraisal of stimuli^34,35^. Moreover, we argue that, while the primary auditory cortex may play a crucial role for the first sensorial processing of the present sound, the centrality of insula and superior temporal pole within the whole-brain network may be essential to predict the forthcoming sounds building the temporal sequence. In conclusion, by investigating the brain connectivity dynamics within the first 220ms after sounds onset, we described the neural mechanisms that are presumably responsible to make temporal information available to human awareness, as suggested by previous investigations that explored the dynamics of human conscious processing^25^.

### Sound encoding, working memory abilities and musical expertise

A further achievement of this study is represented by the modulation of sound encoding brain networks on the basis of participants’ WM skills and musical expertise. First, general WM was associated to centrality of frontal operculum. This result can be seen in light of the several studies that highlighted the fundamental role of frontal and prefrontal cortices for WM tasks^39–41^. Indeed, in our study higher vs lower WM skills participants had a stronger centrality of the most frontal brain region that formed the significant sound encoding brain network.

Remarkably, when considering auditory WM, we did not observe any difference related to frontal operculum. However, we detected a significant centrality of bilateral insula and right temporal middle cortex. Considering that these regions have been shown important for complex processing of auditory stimuli^42,43^, we may speculate that their involvement in an auditory memory task may offer an additional help to encode sounds. While this may happen for best auditory WM scorers, worst participants may rely only on more primitive auditory cortices.

Finally, the higher centrality of left supplementary motor area and right insula for musicians compared to non-musicians is worthy to be mentioned. Since motor learning is a key feature of musical practice, we claim that musicians may recruit also motor areas when encoding sounds. This result could be seen in light of several studies that showed the role of motor brain areas in musicians during music listening^44–46^. Similarly, also insula has been shown more active when musicians compared to non-musicians listened to an early rehearsed and familiar musical piece^47^.

### Conclusions

In conclusion, this study revealed the rapid spatiotemporal dynamics of brain activity and connectivity underlying encoding and prediction of sounds forming a complex temporal sequence. Remarkably, the integration between activity and connectivity provided us with a complete picture of the brain networks involved in this complex cognitive process, networks that the sole brain activity missed to reveal. Notably, our investigation within the first 220 ms after sounds onset allowed us to detect the brain mechanisms that are presumably responsible to make temporal information available to human awareness^25^.

Furthermore, our study highlighted the potential of DFC and phase synchronization analyses to unravel the rapid transition of the brain connectivity patterns from primary auditory cortex to higher order association areas. Indeed, our results suggested that primary auditory cortex centrality would be implicated in the first elaboration of the present sound, while the integration between insula and superior temporal pole with the rest of the brain may play a crucial tole to predict the forthcoming events of the temporal sequence.

Finally, the DFC approach allowed us to detect also the subtle differences among brain regions centrality of participants divided according to their WM abilities and musical expertise.

Taken together, these results remarkably advanced our knowledge of how the brain rapidly encodes and predicts temporal sequence information and provided a first glimpse of the general mechanisms underlying pattern encoding in the human brain.

## Methods

### Participants

Seventy volunteers participated in the study. However, the analyses were carried out on 68 of them since two have been excluded because of technical problems during part of the data acquisition. Thus, the final sample was composed by 68 participants (35 males and 33 females, age range: 18 – 42 years old, mean age: 24.88 ± 4.17 years). Since our experiment involved a well-known piano musical piece, we recruited 23 classical pianists (13 males and 10 females, age range: 18 – 34 years old, mean age: 24.83 ± 4.10 years), 21 non-pianist musicians (10 males and 11 females, age range: 42 – 19 years old, mean age: 24.29 ± 5.02 years) and 24 nonmusicians (12 males and 12 females, age range: 21 – 35 years old; mean age: 25.46 ± 3.48 years).

All experimental procedures complied with the Declaration of Helsinki – Ethical Principles for Medical Research and were approved by the Ethics Committee of the Central Denmark Region (De Videnskabsetiske Komitéer for Region Midtjylland) (Ref 1-10-72-411-17).

### Experimental design and stimuli

Participants’ brain activity was recorded by using MEG. At first, we had a resting state session that has been used later as baseline for evaluating the brain functional connectivity during the task. Participants were required to sit down in the scanner for 10 minutes trying to relax but without closing their eyes. The room was dark and participants were asked to fixate a cross on the screen and not to think about anything in particular. Then, to study the brain dynamics of sound encoding, we asked participants to actively listen to a MIDI homo-rhythmic version of the right-hand part of the entire prelude in C minor BWV 847 by Johann Sebastian Bach, as depicted in **Figure 1A**. Participants were required to try to memorize the prelude as much as possible. For this reason, as well as to collect more data, we played the musical piece four times. The stimuli were designed by using Finale (MakeMusic, Boulder, CO) and then presented through Presentation software (Neurobehavioural Systems, Berkeley, CA). After the MEG recording, in the same or in another day, participants’ brain structural images were collected by magnetic resonance imaging (MRI) exam. Furthermore, participants’ general and auditory WM abilities and musical expertise were assessed. Specifically, their musical expertise was collected using the Gold-MSI questionnaire^48,49^. With regards to general WM we adopted one of the most used psychological tests for assessing cognitive abilities, namely WAIS-IV^50,51^, while for the auditory WM abilities we employed the MET^52^, a newly developed tool that presents couple of complex melodies requiring participants to state whether they are the same or different.

### Data acquisition

We acquired MEG and MRI data in two independent sessions. The MEG data was collected using an Elekta Neuromag TRIUX system (Elekta Neuromag, Helsinki, Finland) equipped with 306 channels. The scanner was located in a magnetically shielded room at the Aarhus University Hospital, Denmark. The data was collected at a sampling rate of 1000 Hz with an analogue filtering of 0.1–330 Hz. Before the exam, we adjusted the sound volume at 50 dB above the participants’ minimum hearing threshold. Furthermore, by using a 3D digitizer (Polhemus Fastrak, Colchester, VT, USA) we recorded the participant’s head shape and the continuous location of four headcoils, with respect to three anatomical landmarks (nasion, and left and right preauricular points). This data was then utilized to ensure a high-quality co-registration of the MEG data with the anatomical structure obtained during the MRI exam.

The location of the headcoils was registered during the whole recording session using a continuous head position identification (cHPI) and therefore we tracked the exact head position within the MEG scanner at each moment. This allowed us to perform an accurate movement correction at a later stage of data analysis.

The MRI data consisted in structural T1. The acquisition parameters for the scan were: voxel size = 1.0 x 1.0 x 1.0 mm (or 1.0 mm^3^); reconstructed matrix size 256×256; TE of 2.96 ms and TR of 5000 ms and a bandwidth of 240 Hz/Px. Each individual T1-weighted MRI scan was then co-registered to the standard Montreal Neurological Institute (MNI) template brain through an affine transformation and referenced to MEG sensors space by employing the Polhemus head shape data and the three fiducial points collected during MEG session.

### Data pre-processing

We conducted Maxfilter^53^ noise reduction on the raw MEG sensor data (204 planar gradiometers and 102 magnetometers) for attenuating the interference that originated outside the scalp by applying signal space separation. Moreover, Maxfilter allowed us to correct for head movement and down-sample the data from 1000 Hz to 250 Hz.

The data was converted into Statistical Parametric Mapping (SPM) format and further processed in Matlab (MathWorks, Natick, Massachusetts, United States of America) by utilizing Oxford Centre for Human Brain Activity Software Library (OSL), a freely available toolbox that relies on a combination of Oxford Centre for fMRI of the Brain Software Library (FSL)^54^, SPM^55^ and Fieldtrip^56^, and in-house-built functions.

The data was high-pass filtered (0.1 Hz threshold) to remove too low frequencies for being originated by the brain. We also used a notch filter (48 - 52 Hz) to control for interference of the electric current. The data was then down-sampled again to 150 Hz and few segments of the data, altered by large artifacts, were discarded after visual inspection. Then, to correct for eyeblinks and heart-beat artifacts, we calculated independent component analysis (ICA) to decompose the original signal in independent components. Then, we individuated and discarded the components that picked up the heart-beat and eyeblink activities and we reconstructed the signal by using only the remaining components^57^. Finally, data was epoched according to the beginning of each of the 605 musical tones of the prelude (pre-stimulus time of 100 ms) and baseline corrected by removing the mean value of the pre-stimulus baseline from the entire trial. Therefore, our trials were represented by the segment of the signal starting with the presentation of each musical tone. The same procedure was carried out also for the resting state. As conceivable, the resting state did not have any external stimulation, therefore we created trials in equal length and number at pseudorandom time-points of the recorded data.

### Event related fields and power spectra analysis

Prior to performing connectivity analysis, as illustrated in **Figure 1B**, we tested the quality of our data by assessing the ERF and especially the N100, a well-known component of the ERF arising 100 – 150 ms after sound stimulation^58^. To this purpose, we averaged together all trials obtained after epoching the data and combined planar gradiometers by mean root square^59^. Then, we calculated a t-test for each MEG gradiometer channel and time-point between the ERF to the sound and the averaged pre-stimulus brain activity. To correct for multiple comparisons, we adopted MCS^60^. Specifically, we reshaped the previously calculated statistics for obtaining, for each time-point, a two-D approximation of the MEG channels layout and we binarized it according to the *p*-values obtained from the previous t-tests (threshold = 1.0e-12). The resulting three-D matrix (*M3*) was therefore composed by 0s when the t-tests were not significant and 1s when they were. Then, we made 1000 permutations of the elements of the original binary matrix *M3*, identified the maximum cluster size of 1s for each permutation and built the distribution of the 1000 maximum cluster sizes. Finally, we considered significant the original clusters that had a size bigger than the 99.9% of the permuted data maximum cluster sizes.

To assess the contribution of each frequency, we estimated the power spectra of the brain signal by employing complex Morlet wavelet transform (from 1 to 40 Hz with 1-Hz intervals)^61^. Then, we calculated a t-test for each time-point within the range: 0.050 – 0.200 seconds and the averaged power spectra of the 100ms pre-stimulus baseline. Emerging *p*-values were binarized according to threshold = 1.0e-18 and then submitted to a two-D MCS. Specifically, we calculated the clusters size of continuous significant values in frequency and time and then made 10000 permutations of the binarized *p*-values. For each permutation we detected the size of the maximum emerging cluster and built a reference distribution with one value for each permutation. Then, we considered significant the original clusters that had a size bigger than the 99.99% of the permuted data maximum cluster sizes.

Thresholds for binarizing the *p*-values matrices were very low since we were comparing the brain activity vs baseline and therefore, as conceivable, the results were highly significant and, to individuate the strongest contribution of MEG channels, time-points and frequencies, it was necessary to select very low thresholds.

### Source reconstruction

As depicted in **Figure 1B**, the brain activity recorded on the scalp by the MEG sensors was reconstructed in source space by adopting an overlapping-spheres forward model and a beamformer approach as inverse model^62^, with an eight-mm grid and both planar gradiometers and magnetometers. The spheres model represented the MNI-co-registered anatomy as a simplified geometric model utilizing a basic set of spherical harmonic volumes^63^. The beamforming used a different set of weights sequentially applied to the source locations for individuating the contribution of each source to the activity recorded by the MEG sensors at each time-point^62,64^.

To assess the brain activity associated to the sound encoding task we submitted the beamformed reconstructed activity to first-level statistical analysis carried out by calculating a GLM for each time-point and at each dipole location^65^. Then, after calculating the absolute value of the reconstructed time-series to avoid sign ambiguity of the neural signal, we conducted group-level analysis, using one-sample t-tests with spatially smoothed variance obtained with a Gaussian kernel (full-width at half-maximum: 50 mm). Finally, to correct for multiple comparisons, a cluster-based permutation test^65^ with 5000 permutations has been calculated on group-level analysis results. Considering an a level = .05, we used a cluster forming threshold *t*-value = 1.7.

### Static functional connectivity

To investigate functional connectivity, we used the source localized data obtained by the beamforming algorithm. Then, this data was constrained into the 90 non-cerebellar regions of the AAL parcellation, a freely and widely-used available template^66^, in line with previous MEG studies^67–69^. This procedure has been carried out for: 2 – 8 Hz, the frequency band characterized by the highest power, according to previous analysis described above. We chose: 2 – 8 Hz instead of: 2 – 5 Hz, as emerging from the power spectra statistics, to have a broader frequency range usually employed in studies on theta waves. Since the length of each musical tone was quite short (36 time-samples with our sampling rate of 150 Hz), to estimate more reliable SFC through Pearson’s correlations, we concatenated and sub-averaged groups of trials. This procedure returned a final time-series matrix *M*, made up by seven concatenated sub-averaged trials, with dimensions: 90 brain regions x 252 time-points. Then, we performed source leakage correction by orthogonalization^70^ and calculated Pearson’s correlations between the envelope of the time-series of each couple of brain areas. This procedure was carried out for both task and resting state (used as baseline) and resulted in two 90 x 90 matrices for each participant, one for the task and one for the baseline. Those two matrices were contrasted by applying Wilcoxon signed-rank test for each couple of brain areas. The resulting *z*-values matrix *Z* was submitted to a degree MCS for assessing which brain area was significantly central within the brain network. In graph theory, the degree of each vertex *v* (here each brain area) of the graph *G* (here the matrix *Z* describing the whole brain connectivity) is given by summing the connection strengths of *v* with the other vertexes of *G*, returning a value of the centrality of each *v* in *G*^71^. In this MCS, we computed the degree of each vertex of *Z*, obtaining a 90 x one vector (*s_t_*). Then, we made 10000 permutations of the elements in the upper triangle of *Z* and we computed a 90 x one vector *d_v,p_* containing the degree of each vertex *v* for each permutation *p*. Combining vectors *d_v,p_* we obtained the distribution of the degrees calculated for each permutation. We considered significant the degrees stored in *s_t_* that were higher than the 99.9% of the degree distribution values calculated by permuting *Z* 10000 times. The threshold of 99.9% derived from simulations of the MCS function with matrices composed by uniformly distributed random values. Setting a 99.9% threshold yielded to a number of false positive nearly equal to zero, while a more common 95% threshold gave rise to few false positives.

### Phase synchronization estimation

To unravel the brain dynamics of the sound encoding, we studied the phase synchronization over time between brain areas for theta band.

By applying Hilbert transform^72^ on the envelope of the reconstructed time-courses (matrix *M* described in the paragraph above) we obtained the analytic signal *S*_(*n_i_,t*)_ expressed by the following equation:

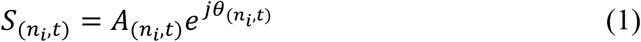

Where *A*_(*n_i_,t*)_ refers to the instantaneous amplitude and *θ*_(*n_i_,t*)_ to the instantaneous phase of the signal for brain region *n_i_* at time *t*. A graphical depiction of Hilbert transform is reported in **Figure 1C**. Then, since matrix *M* was made up by seven concatenated sub-averaged trials, after estimating the instantaneous phase, we discarded the time-samples corresponding to the first and last trials to prevent boundary artefacts introduced by instantaneous phase estimation and we averaged the remaining five, obtaining a 90 brain region instantaneous phases x 36 time-samples matrix *M2*. To estimate the phase synchronization between two brain areas *n_i_* and *n_m_* of the matrix *M2* at time *t*, we calculated the cosine similarity expressed by equation (2)^73^:

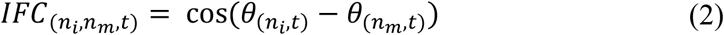

We carried out this procedure for each time-point and each couple of brain areas of the matrix *M2*, obtaining one 90 x 90 symmetric instantaneous functional connectivity (IFC) matrix for each time-point showing the phase synchronization of every couple of brain areas. This procedure was carried out for both task and resting state and is illustrated in **Figure 1D**.

### Brain dynamics of sound encoding

To estimate the instantaneous connectivity matrix *Zt* associated to the task, for each time-point, we contrasted the task IFC matrix at time *t* vs the resting state IFC matrices averaged over time by using Wilcoxon sing-rank test. Then, as described above, an MCS computed on *Zt* assessed the significantly central brain regions within the brain network for each time-point. We refer to this measure as instantaneous brain degree (*IBD_t_*). Moreover, the sum (*sIBD*) over time of *IBD_t_* for the brain area *n* showed us its centrality within the whole-brain network during the whole sound encoding process, as expressed by equation (3):

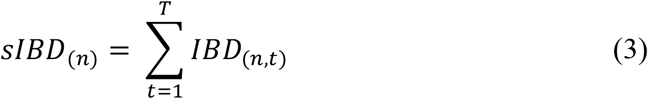

Considering the noise inherent in the data and potentially added by signal processing, we focused our attention towards two main equally-long time-windows (**Figure 1E1**), expressed by the two 90 x one vectors *sIBD*_(*n*)1–110*ms*_ and *sIBD*_(*n*)111–220*ms*_. Then, for each brain region *n* we calculated the vector difference *vtdo*, as expressed by equation (4):

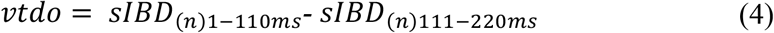

Finally, to assess whether vector *vtdo* contained any significant value, we used MCS. Specifically, we made 10000 permutations of the matrices *IBD*_1–110*ms*_ and *IBD*_111–220*ms*_ and we summed the two matrices over the temporal dimension obtaining two 90 x one vectors, *vto_p_* and *vtv_p_*. Subtracting *vtv_p_* from *vto_p_* we obtained the difference vector *vtd_p_* for the permutation *p*. Performing this procedure for every permutation, we obtained the distribution of the difference vectors. Then, we compared the original differences in vector *vtdo* with the distribution of difference vectors, independently for positive and negative differences, in order to get their associated *p*-values. Finally, we considered significant the brain areas whose *p*-values were lower than:

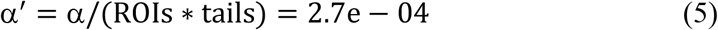

where α corresponds to a level = .05, ROIs are the number of brain regions (90) and *tails* refers to the two tails of the normal distribution of difference vectors created by MCS. In other words, since our hypothesis was that some brain regions were different in the two time-windows, but we did not hypothesize which ones, we looked at the results for each brain region and each direction of the difference (time-windows one > time-windows two and vice versa). Therefore, we had to correct for multiple comparisons by calculating the new threshold α′ expressed by equation (5).

### Brain dynamics of sound encoding and individual differences

In conclusion, as illustrated in **Figure 1E2**, we aimed to assess whether the neural networks underlying sound encoding differed across participants grouped according to general and auditory WM, and musical expertise. To highlight more clearly the differences, we selected for each WM test two groups formed by participants whose scores where at least one SD a part from each other. We adopted this approach since we wanted to include only groups of participants that were clearly differentiated by the tests. With regards to the WAIS test groups the best scorers had a range of 110 – 130, while the worst of 76 – 93 (according to the standardization of the WAIS test one SD corresponds to 15). In relation to MET test, we observed that our best scorers had a range of 43 – 52 while worst scorers of 28 – 36. The mean across all participants was 40.24 with a standard deviation of 6.28. Thus, also in this case the two groups were differentiated by at least one SD. Finally, for musicianship, we considered the 24 non-musicians and the musicians (both pianists and non-pianists) that received a formal musical education for at least 10 years. Those participants were 24. This threshold was set in order to compare people with no musical expertise at all with individuals who engaged in a long-term professional education. Then, as described in the previous section, we focused on the number of times that each brain region was significantly central within the brain, contrasting those values across the two groups by an MCS analogous to the one described above. This procedure was carried out independently for the two WM and the musicianship analyses (since we had three independent tests, we divided the threshold α′ described by equation (5) by three, obtaining a new threshold = 9.2e-05). In **Figure SF1**, to provide full information, brain regions centrality is depicted also for the remaining participants.

## Data availability

The code and multimodal neuroimaging data from the experiment are available upon reasonable request.

## Acknowledgements

We thank Riccardo Proietti, Giulio Carraturo, Mick Holt, Holger Friis for their assistance in the neuroscientific experiment. We also thank the psychologist Tina Birgitte Wisbech Carstensen for her help with the administration of psychological tests and questionnaires.

The Center for Music in the Brain (MIB) is funded by the Danish National Research Foundation (project number DNRF117). Additionally, we thank the Italian section *of Mensa: The* International High IQ Society for the economic support provided to the author Francesco Carlomagno and the University of Bologna for the economic support provided to the author Giulia Donati and the student assistants Riccardo Proietti and Giulio Carraturo.

MLK is supported by the ERC Consolidator Grant: CAREGIVING (n. 615539), Center for Music in the Brain, funded by the Danish National Research Foundation (DNRF117), and Centre for Eudaimonia and Human Flourishing funded by the Pettit and Carlsberg Foundations.

GD is supported by the Spanish Research Project PSI2016-75688-P (AEI/FEDER, EU), by the European Union’s Horizon 2020 Research and Innovation Programme under grant agreements n. 720270 (HBP SGA1) and n. 785907 (HBP SGA2), and by the Catalan AGAUR Programme 2017 SGR 1545.

JC is supported by Portuguese Foundation for Science and Technology CEECIND/03325/2017, Portugal.

## Author contributions

LB, EB, MLK and PV conceived the hypotheses and designed the study. LB, FC, GDO JC, NTH, MLK performed pre-processing and statistical analysis. GDE, EB, MLK, GDO and PV provided essential help to interpret and frame the results within the neuroscientific literature. LB wrote the first draft of the manuscript and, together with FC and MLK, prepared the figures. All the authors contributed to and approved the final version of the manuscript.

## SUPPLEMENTARY MATERIALS

As follows, supplementary materials related to this study and organized as supplementary figures (i) and tables (ii). In the cases when the supplementary tables were too large to be conveniently reported in the current document, they have been reported in Excel files that can be found at the following link: https://www.dropbox.com/sh/z1xdxllcf28ohd1/AACIXDQnoLYjoV7hOaPDmvB7a?dl=0

### SUPPLEMENTARY FIGURES

**Figure SF1.**
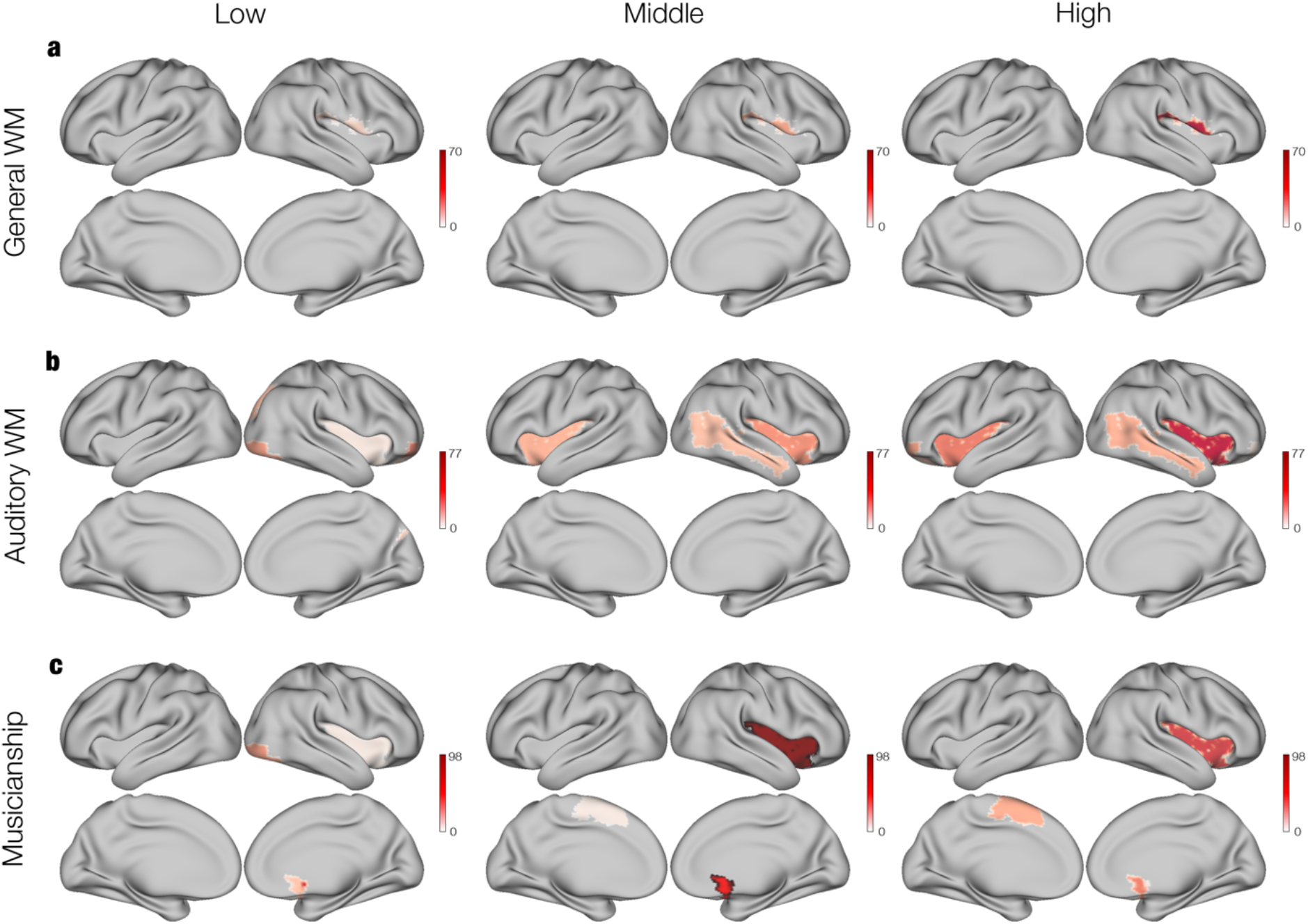
Brain areas centrality according to participants’ WM skills and musical expertise. Brain regions centrality depicted for all participants, divided according to their general and auditory WM skills as well as musical expertise. Best scorers are represented by the right plotted brains, middle scorers by the middle brains and worst performers by the left ones. Colorbars show the amount of time (in ms) when the brain regions were significantly central within the whole-brain network. **a** – Plots related to general WM skills. **b** – Plots linked to auditory WM skills. **c** – Plots connected to musical expertise.

**Figure SF2.**
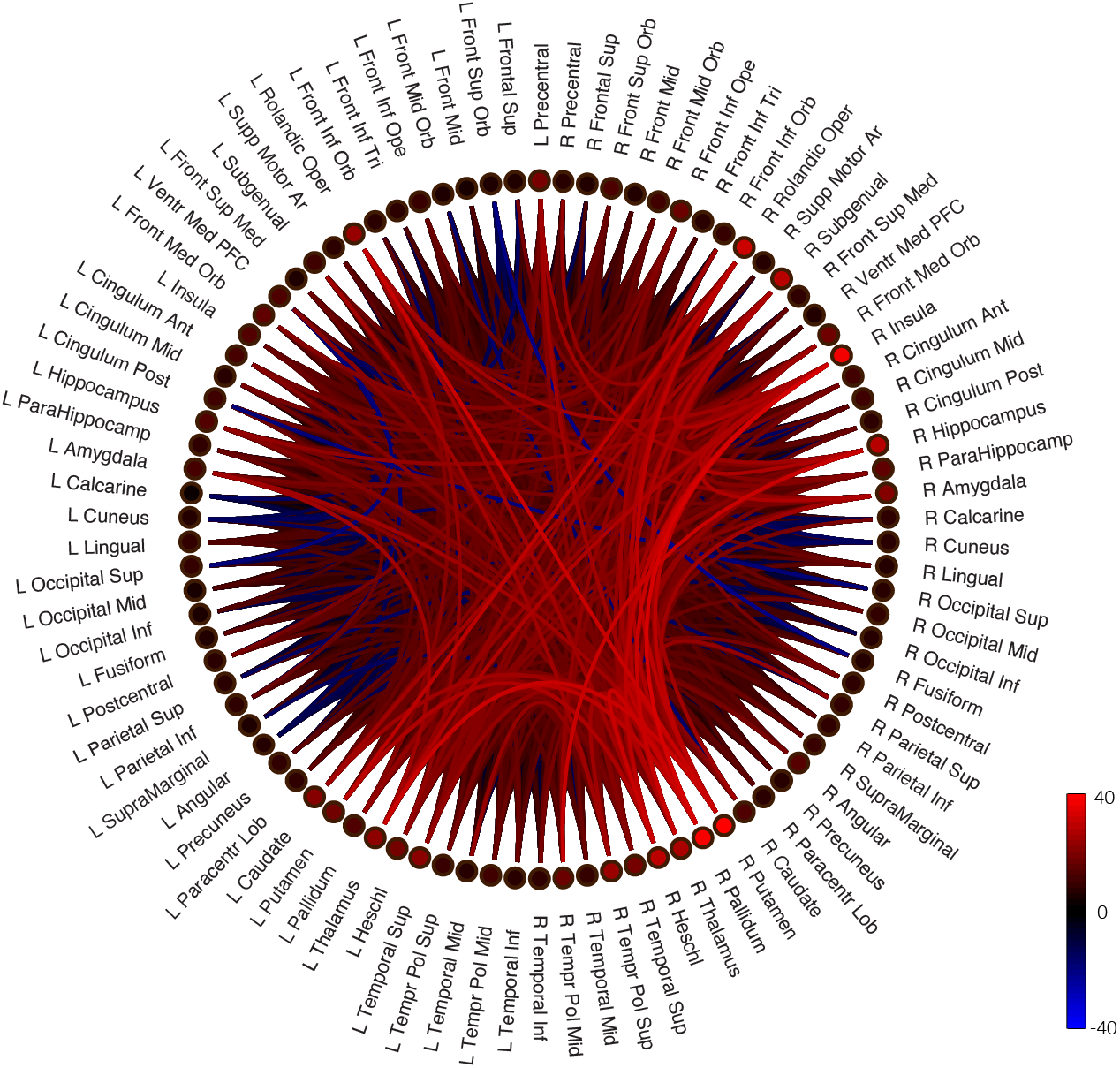
Sound encoding SFC - schemaball. Schemaball representation of the SFC between brain regions concerning the sound encoding task.

**Figure SF3.**
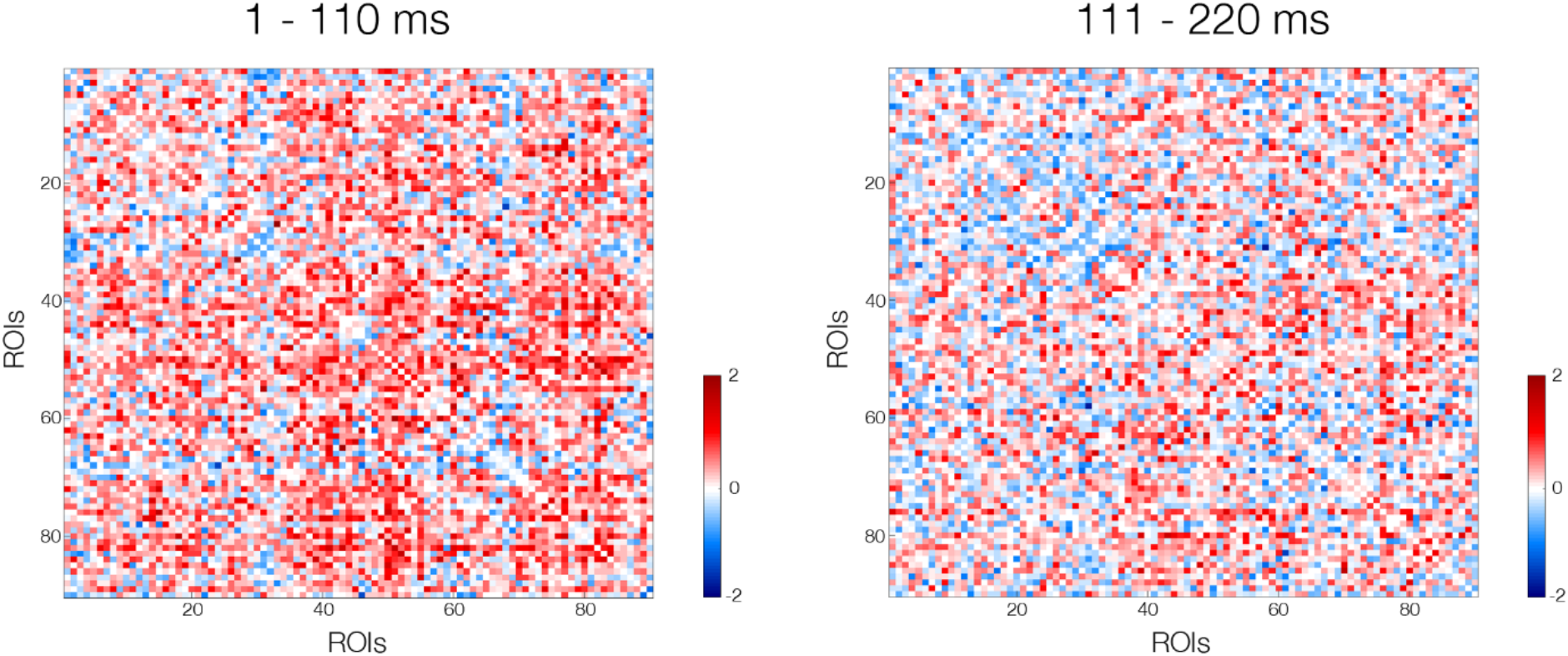
DFC matrices of sound encoding. Matrix representation of the DFC between brain regions during two subsequent time-windows of the sound encoding task.

### SUPPLEMENTARY TABLES

***Table ST1 – Detailed information of significant N100 clusters for MEG sensor data***

Significant clusters of MEG sensors emerged from MCS contrasting Bach’s original vs variation. The excel file depicts those clusters with regards to significant channels and time-windows.

***Table ST2 – Detailed information of significant N100 clusters for MEG source data*** Significant clusters of MEG sources emerged from MCS contrasting Bach’s original vs variation. The excel file depicts those clusters with regards to significant voxels, time-windows and averaged *t*-values for each voxel.

